# Glucose uptake in mammalian cells measured by ICP-MS

**DOI:** 10.1101/2021.10.14.454503

**Authors:** Natalie J. Norman, Joyce Ghali, Tatiana L. Radzyukevich, Judith A. Heiny, Julio Landero-Figueroa

**Author notes:** <u>authors Norman and Ghali contributed equally to this work</u>.

## Abstract

We developed a sensitive, ratiometric method to measure simultaneously ^13^C-labeled glucose and rubidium in biological samples using ICP-MS. The method uses probe-assisted ultra-sonication with water to extract ^13^C-[6C]-labeled-D-glucose and other polar analytes from mammalian tissues. It extracts >80% of the reference value for Rb and >95 % of ^13^C in a CRM spiked with ^13^C-[6C]-labeled-D-glucose in the micro-molar range. Using optimized instrument conditions, the method achieves a stable ^13^C/^12^C signal without spectral interferences. The ^13^C/^12^C signal is independent of sample composition and depends linearly on the concentration of ^13^C-[6C]-labeled-D-glucose in spiked samples. Overall, the method achieves a limit of detection of 10 µM for 6-C-labeled ^13^C glucose in biological tissues. This detection capability for carbon in biological matrices by ICP-MS opens a wider range of applications for ICP-MS in biomedical research. As proof-of-principle, we combined ^13^C detection with the multi-channel capability of ICP-MS to measure glucose and rubidium uptake in the same contracting skeletal muscles. Multi-isotope detection is needed to study many biological processes, including coupled membrane transport. These results demonstrate a capability for carbon detection by ICP-MS that can significantly advance studies of complex biological processes that require multi-isotope detection.

## INTRODUCTION

Glucose is the most abundant carbohydrate in living organisms and is the preferred energy source in mammalian cells. The ability to measure glucose is of great importance in biomedical research and clinical applications, including studies of energy transport and metabolic disorders such as diabetes [1, 2].

The available methods for measuring glucose content or transport in biological samples include enzyme-based assays (glucose oxidase, glucose dehydrogenase, hexokinase et al.) linked to chromogenic reactions or to reactions that generate electron flow [3]; and radio- or fluorescent-labeled glucose used at tracer concentrations [4-7]. These methods are powerful and effective but suffer from difficulties of calibration, consistency among different assays, and potential interference by other components in complex biological samples. Importantly, practical considerations typically limit these assays to detection of a single element or molecule at a time. These methods have been widely used to study glucose uptake into cells by transporters such as GLUT family proteins [8, 9], which transport glucose passively into cells by facilitated diffusion. However, single channel assays provide incomplete information for studies of glucose uptake when it is coupled to the transport of another ion as occurs, for example, with sodium glucose-linked transporters (SGLTs), which import glucose together with Na)[10, 9]. A method that can measure more than one element or molecule simultaneously is needed to fully investigate the mechanisms that underlie coupled glucose transport into cells and tissues.

The goal of this study was to develop a method based on ICP-MS, capable of measuring glucose uptake into live cells while simultaneously measuring other ions that may be co-transported with glucose or whose transport is also stimulated during contraction. The multi-channel capability of ICP-MS is a particular advantage for studies of coupled or secondary transport mechanisms that involve multiple ions species and/or transporters. Using certified a reference material (CRM) bovine liver, we developed a ratiometric method to measure ^13^C-labeled glucose against the large background of natural abundance carbon in biological samples, with negligible interference from other ions present in biological matrices. As proof-of concept, we measured glucose and Rb uptake simultaneously in contracting mouse skeletal muscles. This experimental model was chosen because muscle contraction dramatically stimulates both glucose uptake [11] and NKA transport, which provides the Na gradient that drives many secondary co-transporters.

ICP-MS is an established, sensitive tool for multi-element detection and quantification in a wide range of samples. Technical improvements of the past decade that reduce spectral interferences and increase reproducibility have extended its application to more complex matrices including biological samples. However, biomedical applications of ICP-MS have been largely restricted to detection of inorganic metal and nonmetal ions such as K, Zn, and Fe, P, and S in cells and tissues, including micro-samples [12-14] and single cells [15, 16]. Notably, only a few studies have used ICP-MS to detect C-based materials in the form of micro-plastics[17, 18] but to the time of this work we are not aware of applications in biomedical research. This limited application is due largely to the high carbon background in biological samples, as well as the difficulty in sourcing ^13^C enriched materials. To achieve our goal, we developed an ultra-sonication process to extract C-glucose and other polar targets from mammalian tissues without extracting structural components rich in carbon. We optimized instrument conditions using bovine liver CRM, to obtain a stable ^13^C/^12^C signal with high sensitivity and without spectral interferences. We further validated the method and demonstrated its capabilities for biomedical research by measuring ^13^C-[6C]-labeled-D-glucose and Rb uptake, which are both stimulated by muscle contraction, in the same mouse muscles.

## MATERIALS AND METHODS

### Animals

Adult wild-type female mice (C57BL/6; Jackson Laboratory) at 2-3 months of age were used as a source of tissue. Mice were anesthetized (2.5% Avertin, 17 ml/kg) before tissue extraction and euthanized after tissue removal. All procedures involving animals accorded with the Guide for the Care and Use of Laboratory Animals (National Research Council of the National Academies, USA) and were approved by the University of Cincinnati Institutional Animal Care and Use Committee.

### Chemicals

Chemicals were sourced as follows: Ouabain (Sigma-Aldrich); ^13^C-[6C]-labeled 2-deoxy-D-glucose (Cambridge Isotope Laboratories, Inc., 12 Ci/mmol); [1-^3^H] 2-Deoxy-D-glucose (ViTrax Radiochemicals); Certified Reference Material (CRM) trace metal drinking water (CRM-TMDW, High-Purity Standards, USA; CRM milk powder (CRM-MP, High-Purity Standards, USA); Bovine liver CRM (NIST, 1577b); 999 ± 2 µg/ml in 0.2% (v/v) HNO_3_(CGC1) inorganic carbon standard (Inorganic Ventures, USA). All other chemicals and salts were trace metal grade (Sigma Aldrich, or Thermo Fisher Scientific). Working solutions were prepared using 18 MΩ-cm purity (Milli-Q Academic, EMD Millipore). All vials, pipet tips and materials used were trace metal grade/metal-free or acid washed

### Experimental Solutions

For the mouse muscle experiments, the Equilibration Buffer contained (mM): 118 NaCl (Sigma), 4.7 KCl, 2.5 CaCl_2_, 1.2 MgCl_2_, 1.2 NaH_2_PO_4_, 11 D-glucose, 25 NaHCO_3_; gassed with 95% O_2_, 5% CO_2_; pH 7.4, 32 °C. The Uptake Buffer contained (mM): 118 NaCl, 4.7 KCl, 2.5 CaCl_2_, 1.2 MgCl_2_, 1.2 NaH_2_PO_4_,11 ^13^C-[6C]-D-glucose, 25 NaHCO_3_; and (µM): 200 RbCl. The Wash Buffer contained (mM): 15 Tris-Cl, 2.5 CaCl_2_, 1.2 MgCl_2_, 263 sucrose; pH 7.4, 0–2 °C. All solutions were filtered less than 4 hours before the experiment using 0.22 μm sterile disposable filters (Nalgene Rapid-Flow™, Thermo Fisher Scientific). Solutions were perfused through the chamber at a flow rate of 2 ml/min with the tissue fully submersed. The temperature of the perfusate was monitored by a bath thermistor positioned near the muscle and controlled by an in-line heater. All solutions were stored at 4 °C and used within one week of preparation.

### Tissue dissolution and C extraction for ICP-MS

We evaluated three methods for ^13^C-[6C]-labeled-D-glucose extraction: acid dissolution, water-based mortar-and-pestle, and water-based ultra-sonication. For acid dissolution, muscle samples (8-30 mg) were submersed in a mixture of 100 μL of concentrated sulfuric acid and 100 μL of concentrated hydrochloric acid, and heated on a dry bath for 3 h at 90°C followed by 1 h at 120°C. After the tissue was dissolved, 100 μL of internal standard mix was added to each vial and the samples were brought to a final volume of 10 mL using doubly deionized 18 MΩ water. For the mortar-and-pestle method, approximately 15 mg samples of bovine liver certified reference material (CRM) were weighed and transferred to a mortar. 3 ml of 18 MΩ water was added and the mixture was ground manually using a pestle for 3-5 minutes. The solutions were then centrifuged for 7 minutes at 450 g. The supernatant was transferred to a 0.45 µm spin filter and centrifuged for an additional 5 minutes at 7,000 g. The sonication protocol was optimized as follows: Approximately 15-20 mg of bovine liver CRM was weighed out in a 15 ml metal free tubes to which 3.0 mL of 18 MΩ water was added. The mixture was first vortexed for approximately 5 seconds then ultra-sonicated with a 3 mm x 100 mm sonication probe programmed to deliver 2 s pulses with a 3 s rest at 30% amplitude (37.5 watts) for a total time of one minute. Samples were then split into two equal volume solutions into 1.5 ml centrifuge tubes and centrifuged for 5 minutes at 13,000 g. Next, the supernatants were transferred to a different vial, acidified with 75 µl of concentrated HCl and kept at -20 °C overnight. The samples were then transferred to 0.5 ml, 0.45 µm spin filters and centrifuged for 10 minutes at 9,000 g. Sample solutions were then pooled to obtain a homogenous mixture used to perform the ^13^C-[6C]-D-glucose and magnesium spiking.

For the analysis of mouse muscles, the sonication method was used with cryo-crushing as additional sample preparation step. For this a Cellcrusher tissue pulverizer (Schull, Co. Cork, Ireland) was used. The provided metal spoons and disassembled tissue crusher components were placed in a medium sized foam cooler which was subsequently filled with liquid nitrogen. Tools were chilled for two minutes. The tissues were submerged in liquid nitrogen for two minutes and placed inside the crusher. To pulverize the tissue, the tissue crusher pestle was gently lowered into base using pliers and struck three times using a rubber mallet. Lastly, the tissue crusher pestle was lifted using pliers and chilled metal spoons were used to transfer pulverized tissue on tissue crusher components into a 5 mL metal free Eppendorf tube. The volume of water used for the sonication was 1.5 ml for the EDL and 3 ml for the TA tissues. The sonication and filtration steps were the same as the ones used for the CRM.

### Measurement of rubidium and ^13^C-labeled glucose by ICP-MS

Quantification of the elements of interest by ICP-MS was accomplished using an Agilent 7500ce instrument equipped with a collision cell, a Cetac ASX-500 series auto sampler connected to a micromist nebulizer (Glass Expansion) using 0.25 mm ID PTFE tubing into a double pass Scott-type chilled spray chamber. The torch was a standard 2.5 mm insert quartz torch with platinum shield torch. The cones used for the interface were nickel sample and skimmer cones with a CE lens stack. The instrument was operated in multi-tune isotope analysis mode. Helium mode used for polyatomic interference removal was tuned daily with 1 ppb of Ce, Li, Co, Y and Tl. The no-gas mode was tuned daily with a 1 ppm ^13^C solution in the form of ^13^C-[6C]-D-glucose against a 1 ppm natural abundance carbon in the form of glucose.

### Spiking of ^13^C-[6C]-D-glucose in biological tissues

In order to evaluate the effect of endogenous carbon background on the ^13^C spiking recovery, a portion of the CRM extract was transferred to a different metal-free tube and diluted 2 times with 18 MΩ water by mass before adding the ^13^C-[6C]-labeled-D-glucose spiking solution, while the original extract was spiked without further dilution. For the spiking experiments, 1,350 µL solutions of both the original and diluted CRM solutions were spiked with varying volumes of a 10 ppm ^13^C working standard and diluted with LC-MS grade water to a final volume of 1,500 µL to obtain two sets of CRM solutions. The spiking experiment was then carried out under two levels of carbon background, at the 0.083, 0.165, 0.413 and 0.826 ppm based on ^13^C.

### Carbon Calibration Curves

A 100 ppm working standard was prepared using 999 ± 2 µg/mL in 0.2% (v/v) HNO_3_(CGC1) inorganic carbon standard (Inorganic Ventures, USA), and used to make 0, 0.5, 1, 5, 10, 25, and 50 ppm water-based calibration standards. Using ^13^C-[6C]-labeled-D-glucose (Cambridge Isotope Laboratories, Inc.), 117.785 and 10 ppm ^13^C primary and working standards respectively were prepared and used to make 0, 0.2, 0.4, 0.8, 4, 8 and 20 ppm ^13^C water-based calibration standards.

### Instrumental considerations for reliable carbon isotopic analysis

In order to achieve a reliable carbon signal from the ICP-MS in the intended samples, the following steps were taken: i) The spray chamber, torch and torch connector were cleaned by soaking in a solution of 0.1% triton X-100 for 10 minutes, subsequently rinsed with 18 MΩ water, followed by an overnight soaking in 18 MΩ water. The cleaned set was used exclusively for this analysis. ii) The detector was forced to acquire the signals for both ^12^C and ^13^C in analogue mode, under isotope analysis mode in order to avoid any pulse/analogue factor variations during or between runs. iii) The background signal and the ^13^C/^12^C ratio were pre-monitored and adjusted in the tune window; for this, the ^13^C signal was adjusted between 3×10^4^ -5×10^4^ CPS; for a ^12^C signal of 2.3×10^6^ – 4.0×10^6^ CPS. Values above this range would compromise the response of the detector and, in our experience, are most likely due to contamination of the interface. iv) The stability of the ^13^C/^12^C ratio at different carbon concentrations is sensitive to the ICP-MS lens voltages; for this reason, the lenses were adjusted daily by comparing a 1 ppm to a 50 ppm inorganic carbon standard for a ratio difference below 3%. After the stability of the natural abundance carbon ratio was ensured, a 0.5 ppm ^13^C in the form of ^13^C-[6C]-labeled-D-glucose was used to ensure a proper response in the form of an increased ^13^C/^12^C ratio. v) The rinsing of the sample introduction system is critical for a reliable and stable signal. For this method, rinsing with 18 MΩ water was adopted and, in order to optimize the rinsing time, while keeping the sample consumption to a minimum, manual acquisition was used. The signal pre-monitoring option was set for 25 s after the sample reached the nebulizer. In this way, the acquisition can be manually started or aborted based on the stability and magnitude of the carbon isotope signals. Once optimized (typical values of 10-15 s were observed in normal samples), the time can be set for the use of the auto sampler and the intelligent rinse between samples can ensure that the high concentration standards are rinsed properly.

The instrument used in this study was a quadrupole based ICP-MS and for this, the accuracy of the isotopic ratio is not enough for *de-novo* isotopic distribution studies. Nevertheless, the precision of the instrumental ^13^C/^12^C ratio can be sustained below 3% for a 6h analysis, with a consistent return to the initial value after rinsing with 18 MΩ water. In order to achieve the best instrumental ^13^C/^12^C ratio, an un-treated or un-spiked sample was extracted and analyzed. A summary of typical instrument tune parameters is given in **Table 1**.

**Table 1.** Instrument tune parameters for the ICP-MS quantifications

| Parameter | Tune mode |  |
| --- | --- | --- |
|  | No gas | He gas |
| Forward power | 1600W | 1600W |
| Nebulizer gas flow | 1 L/min | 1 L/min |
| Extract 1 | 3V | 3V |
| Extract 2 | -140V | -140V |
| He gas flow | 0mL/min | 4 mL/min |
| OctP bias | -8V | -18V |
| OctP RF | 140 | 140V |
| Energy discrimination □ | 8.7mV | 8.7mV |
| Isotopes Monitored and Integration Times | $^{12}\text{C}$ & $^{13}\text{C}$: 1<br>$^{5}\text{Sc}$, $^{89}\text{Y}$, $^{103}\text{Rh}$: 0.1 | $^{31}\text{P}$: 0.25<br>$^{31}\text{P}$, $^{45}\text{Sc}$, $^{85}\text{Rb}$, $^{89}\text{Y}$, $^{105}\text{Pd}$: 0.1 |

### Measurement of ^13^C-[6C]-labeled-D-glucose and Rb uptake by EDL muscles

Two extensor digitorum longus (EDL) and two tibialis anterior (TA) muscles were taken from each mouse. The TA muscle was used to obtain the endogenous, basal Rb content of untreated muscles as well as the experimental ^13^C/^12^C ratio under no ^13^C exposure. The endogenous Rb concentration varied 5-10% in different animals, but was highly consistent (<2%) in different muscles from the same animal, as reported [19]. The endogenous Rb concentration of untreated mouse muscles was in the range of reference values for the Rb content of CRM bovine skeletal muscle (NIST RM 8414). The TA muscles were untreated, weighed, placed in an acid-washed vial, stored at 4 °C, and assayed by ICP-MS alongside the EDL samples. To measure glucose and Rb uptake, an EDL muscle was placed in a chamber perfused with Equilibration Buffer at 32 °C and positioned between parallel platinum plate electrodes. One tendon was fixed and the other tendon was attached to a force transducer. The muscle was stimulated with brief pulses (0.5 ms duration) to find and set L_0_, the length at which the muscle produces peak twitch force. Thereafter, the muscle was perfused for 15 minutes at 32 °C in Equilibration Buffer. The muscle was then incubated for 5 min at 32 °C in Uptake Buffer containing 11 mM ^13^C-labeled glucose and 200 µM RbCl, which was used as a tracer for K transport by the NKA. During the uptake period, the test muscle was stimulated electrically to produce repetitive tetanic contractions (brief pulses applied at 90 Hz for 10 seconds, repeated once per minute for 5 min). The contralateral muscle from the same mouse served as control and was subjected to the same protocol but without stimulation. After the uptake period, the muscle was perfused immediately without ^13^C-labeled glucose, K, Rb, or Na wash Buffer at 0 °C, then removed from the chamber and washed in 10 mL of Wash Buffer for 5 minutes at 0 °C, repeated 4 times with shaking. The wash procedure stopped enzyme cycling and removed excess cations from the muscle extracellular spaces. After washing, the muscle was gently blotted, weighed on an analytical balance, placed in an acid-washed glass digestion vial with Teflon-lined cap, and stored at 4 °C until taken for measurement of the ^13^C and Rb content by ICP-MS.

### Measurement of ^3^H-glucose Uptake by EDL muscles

In order to validate our developed method, the glucose uptake measured in our muscle model by ICP-MS was compared with glucose uptake measured using a radio tracer assay in the form of ^3^H-2-deoxy glucose. Essentially the same protocol described above was used to measure ^3^H-2-deoxy glucose uptake, except that the uptake buffer contained 11 mM unlabeled glucose and a tracer amount of ^3^H-2-deoxy-D-glucose. After washing, the muscles were gently blotted, weighed on an analytical balance, and placed in a scintillation vial containing 250 µL formic acid and 100 µL hydrochloric acid. The vials were heated overnight in a water bath at 50 °C to disolve the muscles. Following dissolution, 3 mL of scintillation fluid was added to the vials and the vials were placed on a shaker for 10 min before counting. An aliquot of Uptake Buffer was taken in each experiment to measure ^3^H activity for calibration of glucose uptake.

### Calculation of glucose and rubidium uptake rates from ICP-MS data

The concentration of Rb in the tissue (in ng/mL) was obtained using the standard calibration curve equation generated by plotting a range of Rb concentrations and their respective CPS. The uptake rate of Rb (in nM/g tissue-min) is obtained by multiplying the Rb concentration by the following dimensional factors:

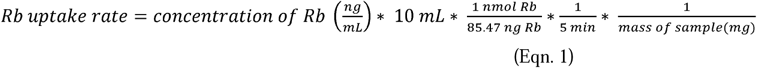

Isotopic abundances obtained with quadrupole based ICP-MS instruments are known to deviate from the natural abundance ones, especially for very light or very heavy elements, as a result of instrumental mass bias. For this reason, it is necessary to experimentally determine the naturally occurring ^13^C/^12^C ratio in the instrumental conditions for each analysis day. This was accomplished by analyzing the untreated TA muscle, which was never been exposed to ^13^C-[6C]-labeled-D-glucose and provided a reference for the naturally occurring ^13^C/^12^C ratio in the EDL muscle from the same animal. This ratio was used to calculate the natural content of ^13^C, and subtracted from the values measured in the treated muscles to obtain the CPS of ^13^C from the labeled glucose using the following equation:

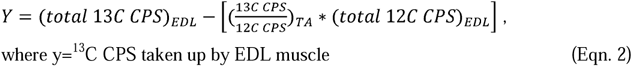

Subsequently, to obtain the concentration of ^13^C taken up by the treated muscle, the calculated extra ^13^C CPS (compared with the predicted ones from the ^12^C signal * ^13^C/^12^C in the TA) was input into the standard calibration curve equation generated using a commercial C standard. The calibration curves were constructed using the natural abundance of each isotope and not just the nominal concentration. This resulted in calibrations based on the individual content of each isotope. The signal for ^12^C and ^13^C was used to calculate the amount of ^13^C-[6C]-labeled-D-glucose taken up using the equation:

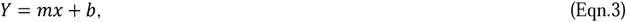

where x= ^13^C CPS taken up by the EDL muscle, y = the concentration of ^13^C in ng/mL obtained from the calibration curve of the ^13^C concentration adjusted for the isotopic distribution of carbon, m=sensitivity, and b = y-intercept.

Finally, the uptake rate of ^13^C-[6C]-labeled-D-glucose (in nmol glucose/ (g-tissue-min)) was obtained by multiplying the concentration of ^13^C (in ng/mL) by the following dimensional factors:

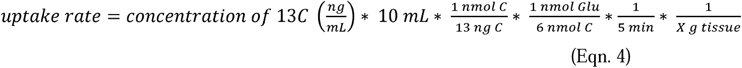

### Data analysis and statistics

Sigma Plot 14 and Origin 2018 (OriginLab Corp.) were used for statistical analyses. Significant differences between means of normally distributed groups were evaluated by Student’s T-Test.

## RESULTS

### Glucose and rubidium extraction from mammalian tissues

The target analytes in this study were ^13^C-[6C]-labeled-D-glucose and ionic rubidium, which are both dissolved in the cytosol of mammalian cells. We optimized Rb extraction using 18 MΩ water as extractant and bovine liver CRM as test matrix. Bovine liver was used to develop the protocol because it was the closest available reference standard with a Rb concentration and sample composition similar to mammalian skeletal muscle. The concentration of Rb (in ppb) extracted was 9,686.6 (± 2,477.8, n=4) for mortar & pestle, 11,130.3 (± 4,471, n= 4) with ultra sonication probe, and 13,566.8 (± 363.8, n=4) with acid dissolution. These represent yields of 70.7, 81.2, and 99.0 % of the CRM reference value (13,700 ppb ± 1,100). Although acid dissolution gave a slightly higher yield, it was rejected because inorganic carbon (charcoal) formed as a by-product and interfered with target signals by absorbing analytes in solution. Consequently, it was necessary to perform filtration on the acid-digested samples as soon as they reached room temperature, to avoid time-dependent decay of the Rb signal (data not shown). The extraction efficiency of ^13^C-[6C]-labeled-D-glucose and Rb are the same between the two methods; however, due to the reduction of the natural abundance carbon background, the sonication protocol was chosen for the ^13^C spiking and Mg interference measurements.

### Carbon analysis by ICP-MS, measurement of background C under different conditions and evaluation of its impact on ^13^C quantification

For all calibrations in this work, the natural abundance of each isotope was taken into account in order to obtain the isotopic sensitivity and not the sensitivity obtained from the nominal mass. In order to obtain a good performance of this method for the intended application, the sensitivity of ^12^C and ^13^C should be similar, and more importantly, stable within the intended working range regardless of the carbon source and background concentration of endogenous carbon.

We started by generating a calibration curve for ^12^C and ^13^C in 18 MΩ water, by using both, an inorganic carbon standard and ^13^C-[6C]-labeled-D-glucose. The calibration range was from 0.5 – 50 ppm based on total carbon; the calibrations based on the isotopic abundance of ^12^C and ^13^C can be seen in figure 1. The slope for ^13^C from the inorganic standard and the ^13^C-[6C]-labeled-D-glucose were not different. From these calibrations, it is evident that the instrument response is positively biased to the ^13^C isotope, which consistently resulted in a greater slope than ^12^C, in the 15-25% range. This highlights the need for an experimental ^13^C/^12^C ratio measurement per running session in untreated samples.

**Figure 1.**
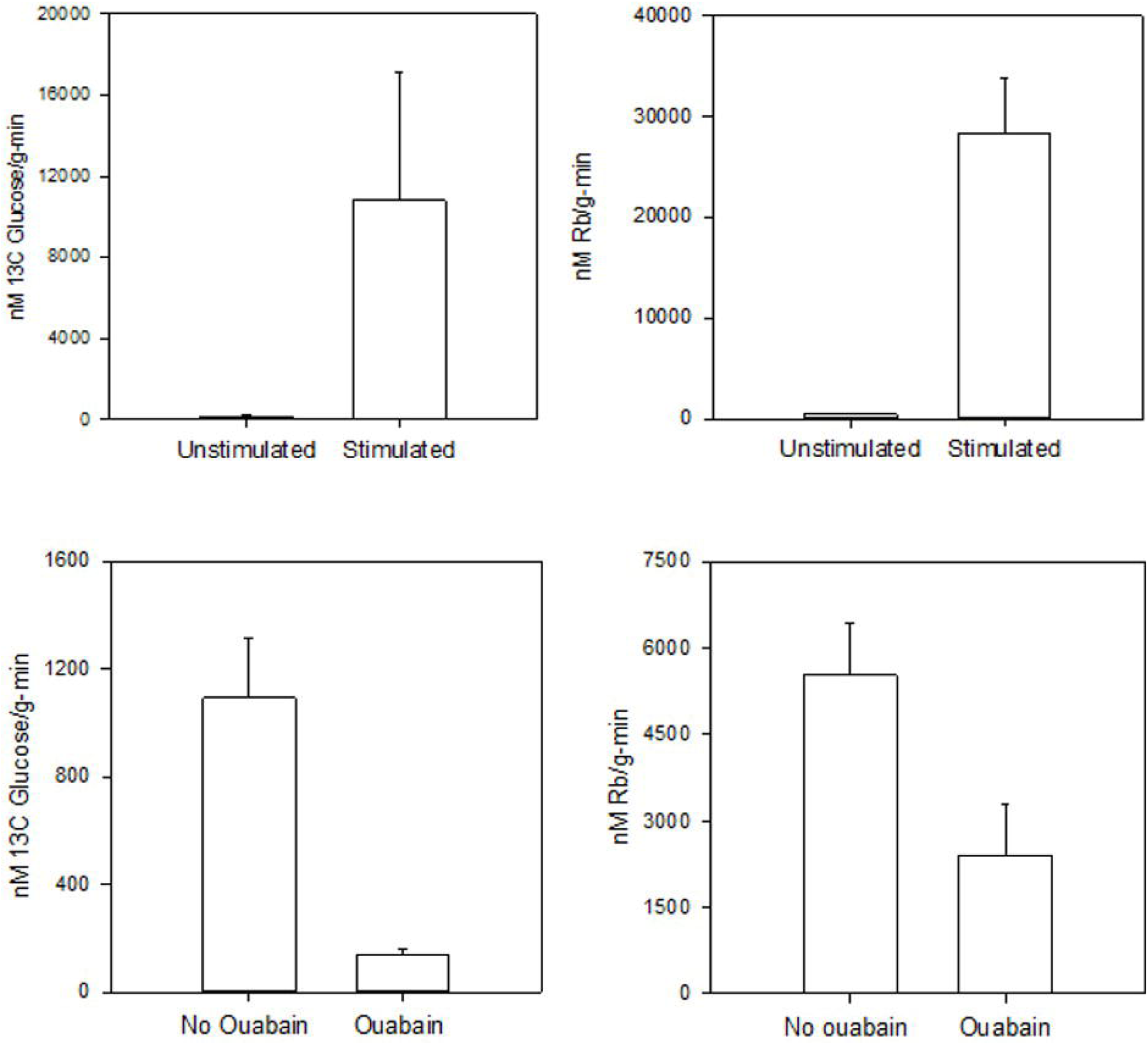
Calibration curves of natural abundance inorganic (a) ^13^C and (b) ^12^C compared to one (c) ^13^C generated with ^13^C-[6C]-labeled-D-glucose. Each point reflects the average of 3 technical replicates with the error bars represent the SD.

Another important parameter from these calibrations is the blank equivalent concentration (BEC), which is the C content in the blank (18 MΩ water in this case). The BEC for ^13^C was ≈ 0.05 ppm while the ^12^C was 4.5 ppm as seen in Supplemental Figure 1. Given that the ability to distinguish the ^13^C signal coming from the ^13^C-[6C]-labeled-D-glucose is limited by the variation of the background signal, and that the absolute CPS variation of the background is a function of the total C signal, minimizing the C background is necessary and important for improving the detection capabilities of the method. The C background in 18 MΩ water (≈ 4.5 ppm) comes primarily from dissolved CO_2_, which is a non-polar molecule and therefore has low solubility in pure water, and from the carbonic acid/bicarbonate forms that result from the hydration of CO_2_. Extensive bubbling for 3 minutes with an inert gas, helium in our case, at approx. 1 L min^-1^, decreased the background signal by about 22% in the 18 MΩ water blank, yet it only decreased the background signal of the extracted control tissues by around 1%. This decrease in the background came at the cost of time and instrumental modifications and it introduced variability if the time of bubbling and analysis was not consistent. This prompted us to discard this marginal improvement to keep the run time shorter and ensure stability of the ICP-MS signal.

The next step in the validation process was to ensure a comparable ^13^C sensitivity between the inorganic carbon standard proposed for use and the enriched ^13^C-[6C]-labeled-D-glucose standards, in both 18 MΩ water and in the extracted tissues at different carbon background concentrations. Figure 2 shows the calibrations obtained for ^13^C from **i)** inorganic carbon standard in 18 MΩ water, **ii)** ^13^C-[6C]-labeled-D-glucose in 18 MΩ water, **iii)** ^13^C-[6C]-labeled-D-glucose in a concentrated CRM extract (15 mg dry mass in 3 ml of 18 MΩ water) and **iv)** ^13^C-[6C]-labeled-D-glucose in a diluted CRM extract (15 mg dry mass in 6 ml of 18 MΩ water). The observed variation in sensitivity from technical and biological replicates were below 5% for ^13^C and ^12^C. The use of the isotopic abundance concentration of ^13^C in the standards translated into a first calibration point being 0.005 ppm, well within the margin of our calculated instrumental LOD for ^13^C (see below). The displacement of the regression lines to higher BECs with the same slope when a spiked tissue is analyzed can be used to quantify the extracted background of endogenous carbon (proteins, other metabolites and soluble bio-molecules in general). It is also an important parameter for further method optimizations and to ensure that the studied tissues under test conditions (electrical stimulation in this study) are within the desired range of background. For our samples, the endogenous carbon background was in the 1.6-3.5 ppm range. It is important to highlight that during this optimization of background range, the important parameter to track is the stability of the ^13^C/^12^C ratio. In our hands, the ratio was stable from 1 to 50 ppm at < 3% for contracted muscle samples, and < 1% in the control tissues with a similar endogenous carbon background (1-2 ppm). The spike recoveries at the concentrated and diluted CRM extracted are illustrated in figure 3a. From this, it is clear that the studied C backgrounds do not impact the ability to quantify a 0.1 ppm spike of ^13^C.

**Figure 2.**
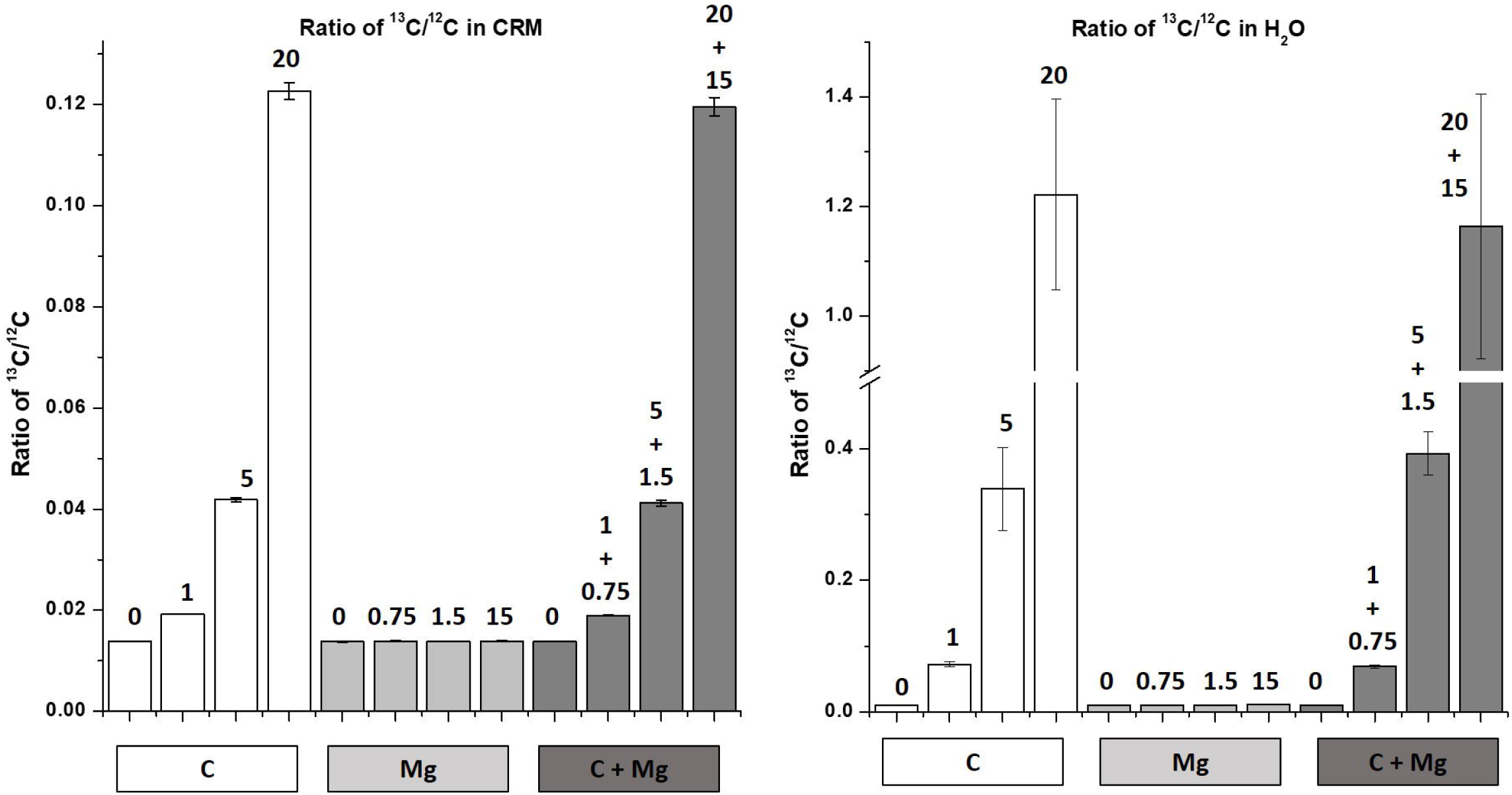
Calibration curves obtained for ^13^C and ^12^C from i) inorganic carbon standard in 18 MΩ water, ii) glucose in 18 MΩ water, iii) ^13^C-[6C]-labeled-D-glucose in 18 MΩ water, iv) ^13^C-[6C]-labeled-D-glucose in a concentrated CRM extract (15 mg dry mass in 3 ml of 18 MΩ water) and v) ^13^C-[6C]-labeled-D-glucose in a diluted CRM extract (15 mg dry mass in 6 ml of 18 MΩ water). Each point reflects the average of 3 technical replicates with the error bars represent the SD.

**Figure 3.**
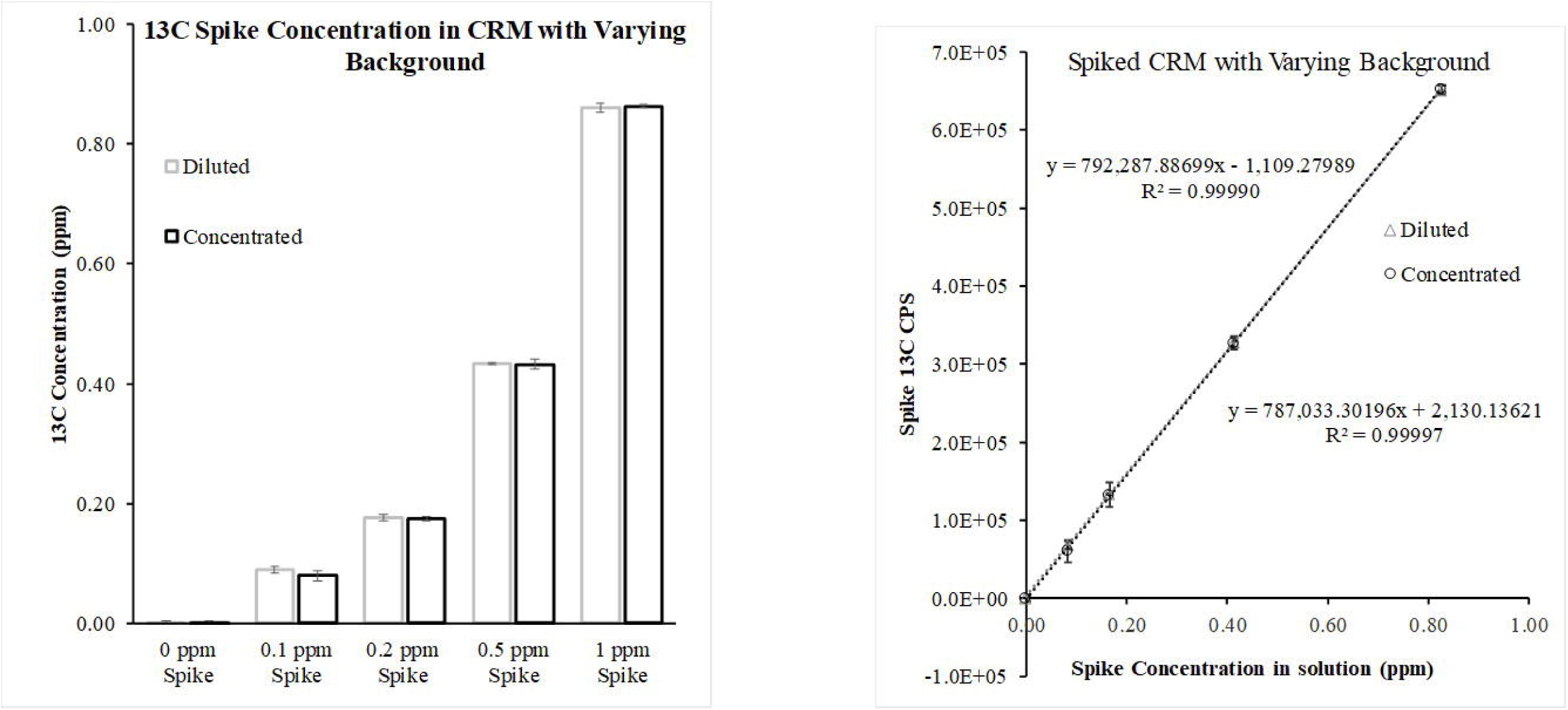
(a)^13^C concentration quantified from four spike levels in bovine liver CRM extracts at two different matrix dilutions, diluted represents 2x lower carbon background. (b) Calibration based on the calculated ^13^C-exogenous content from the spike experiments. The two calibrations correspond to the diluted and concentrated CRM extract, each point represents 3 technical replicates and the error bars the SD.

The use of a standard addition method for the total ^13^C was not an option in this methods given that the total ^13^C concentration in extracts of real samples would be too variable. Instead of the standard addition method, the proposed calculation of the extra-^13^C CPS shows no effect of the carbon background, and when plotted against the spiked concentration (figure 3b), it shows a linear behavior with the same slope (<5% difference, n=3) as the ones observed in the inorganic C and ^13^C-[6C]-labeled-D-glucose standards as seen in Figure 2.

### Mg in mammalian tissue does not interfere with carbon detection

Because mammalian tissues contain Mg at millimolar concentrations, it was important to determine whether the presence of Mg in our samples interferes with detection of carbon by forming the M^2+^ ions ^24^Mg^2+^ and ^26^Mg^2+^. Although the second ionization potential of Mg is only 0.725 eV lower than the first ionization of Ar (15.035 eV for Mg. 15.76 eV for Ar) the concentration of Mg in the samples prompted us to evaluate the remote possibility of these interferences being formed. To address this question, we measured C and Mg in samples of CRM spiked with 0, 1, 5, or 20 ppm of ^13^C only, or with 0, 0.75, 1.5, 15 ppm Mg only, or with both ^13^C and Mg (Figure 4). Spike concentrations for ^13^C were based on the range of concentrations of glucose in muscle diluted by our typical 150x dilution factor, which was approximately 5 ppm based on carbon. Spike concentrations for Mg (ppm) were determined from the Mg concentration in mammalian tissues (0.3 – 2 mM), after factoring in our dilution factors (15 mg of CRM in 3 ml of 18 MΩ water and a subsequent 2x dilution factor). This yielded 1.5 ppm Mg as a starting spike concentration for bovine liver CRM.

**Figure 4.**
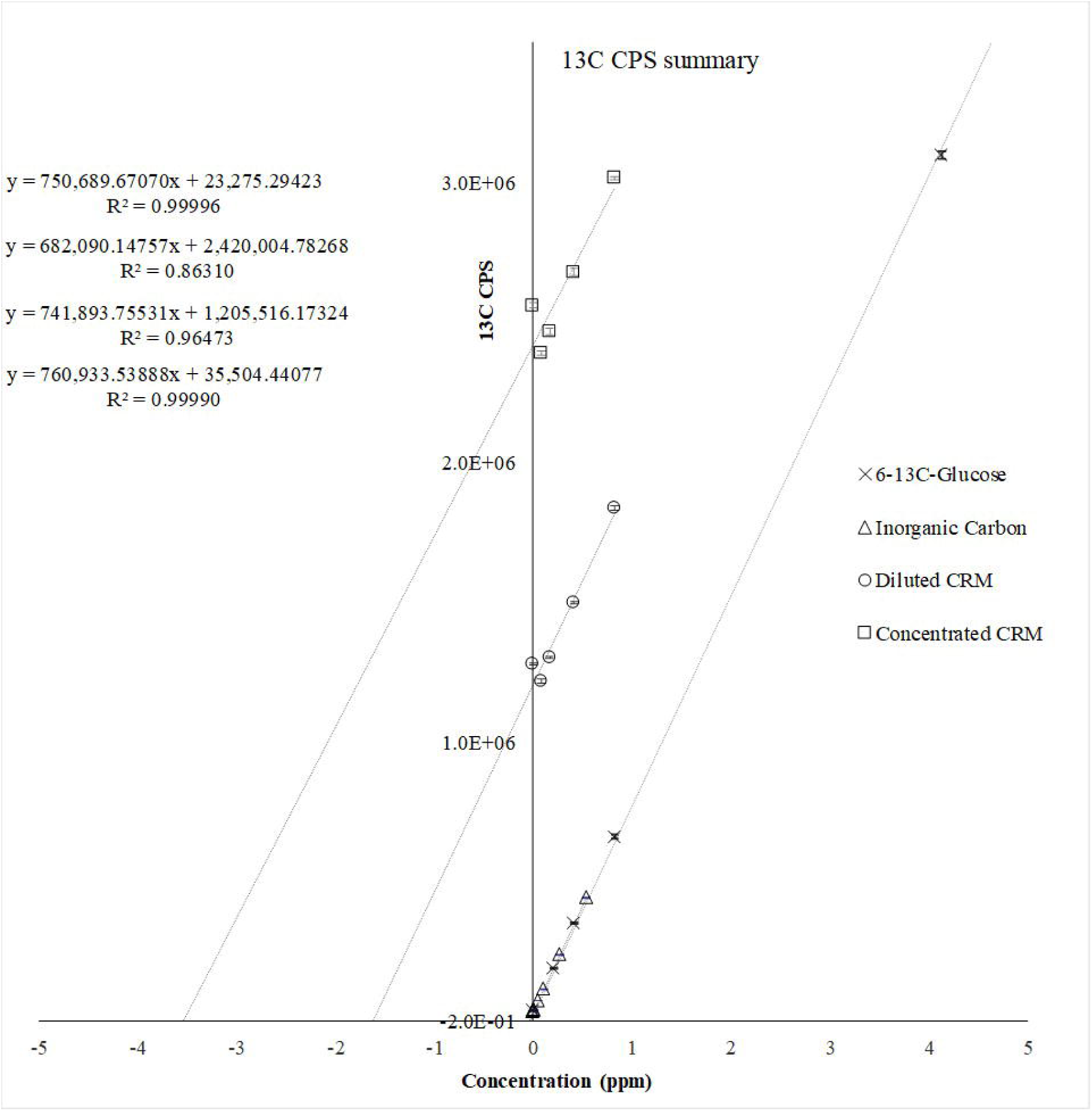
^13^C/^12^C ratios for (a) three spike levels of carbon, magnesium and carbon + magnesium in bovine liver CRM extracts and water spiking experiments. The numbers on top of each bar represent the spiking level of carbon, magnesium or carbon + magnesium. Error bars represent the standard deviation from 3 experimental replicates.

**Figure 5.**
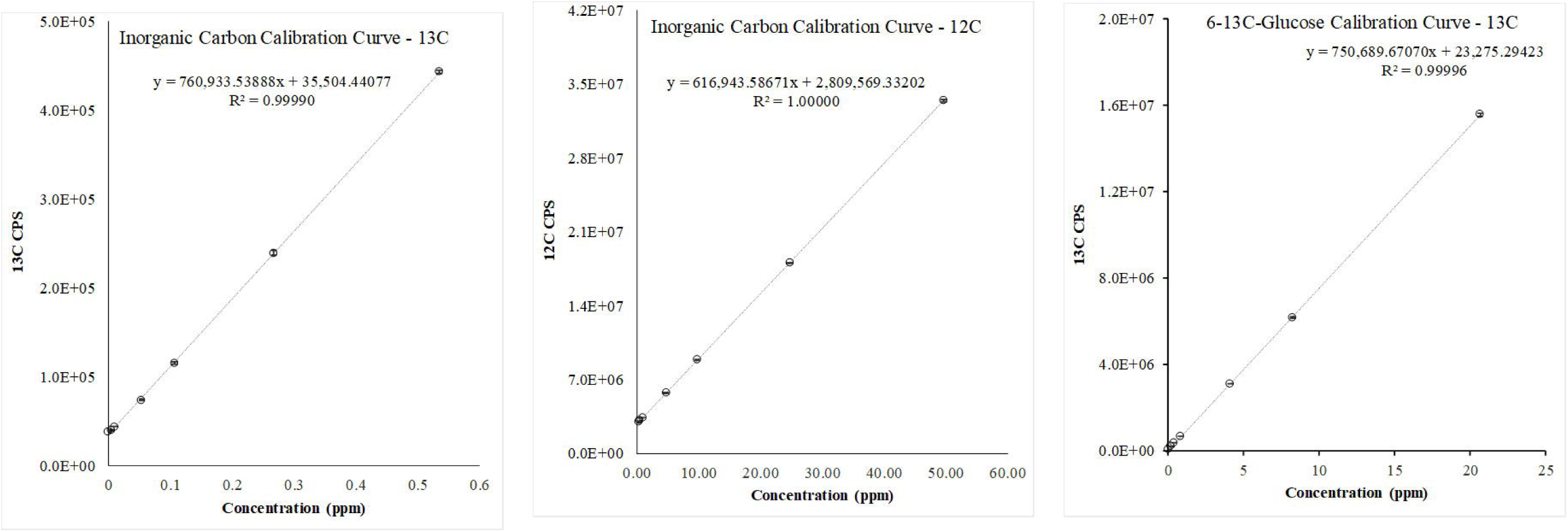
Glucose uptake and Na,K-ATPase activity during muscle contraction. A) ^13^C-[6C]-labeled-D-glucose uptake rate in quiescent and contracting EDL muscles. B) Rb uptake rate measured in the same muscles. An isolated mouse EDL muscle was incubated with 11 mM ^13^C-[6C]-2-deoxy-D-glucose and 200 µM tracer Rb and either left quiescent or stimulated to produce repetitive contractions using conditions that maintain force and do not produce fatigue (10 sec, 90 Hz tetani repeated once per min for 5 min), as described in Methods and Supplemental Fig. 1. The amount of glucose taken up during the contraction period was measured by ICP-MS using the ^13^C/ ^12^C ratio and converted to glucose uptake rate using Eqn. 1. The rate of glucose uptake in quiescent muscles was below detection limits. Uptake rates for Rb were computed after scaling by the ratio of Rb to K in the uptake buffer. Each comparison used paired test and contralateral muscles from n= 3 – 4 animals. C & D) rates of glucose and Rb uptake in stimulated EDL muscles in the absence and presence of 0.75 µM ouabain. n= 4 animals. Bars show group means ± SD/SEM * indicates statistically significant difference at P < 0.05.

The instrument ^13^C/^12^C ratios for un-spiked samples were stable independently of the matrix. This observation suggests no interference of Mg with ^13^C and ^12^C signal, even at maximum applied power of 1600 W. The ^13^C/^12^C ratio scaled linearly with ^13^C concentration. It was independent of Mg at all concentrations, and identical to the C signal in water. The stability of the ^13^C signal across samples spiked with different concentrations of Mg confirms that there are no measurable interferences from ^13^C in the assay. The ratio of ^13^C/^12^C in samples spiked with both ^13^C and Mg (dark grey) were identical to the ^13^C/^12^C ratio in samples spiked with the same concentrations of ^13^C alone (white). These measurements demonstrate that the presence of Mg at physiological concentrations does not interfere with detection of C or Rb in biological samples. The Rb signal (supplemental figure 2) was identical at all concentrations of ^13^C and/or Mg, a result that also confirms the efficiency and consistency of the extraction method.

The mass spectra of the CRM samples spiked with 1.5 ppm Mg and/or 5 ppm ^13^C (Supplemental Figure 3) further confirms this finding. Mg has three stable isotopes with masses 24, 25, and 26. The second ionization potential of Mg (15.035 eV) is slightly below that of the first ionization potential of the Ar (15.760 eV) plasma and near the first ionization potentials of ^12^C & ^13^C (11.266 eV). Consequently, if a small amount of ^24^Mg and ^26^Mg were to form M^2+^ ions, the resulting signals at m/z 12 and 13 would interfere with the carbon signals. The absence of a peak at the 12.5 m/z mark at shown in Supplemental Figure 3 indicates the absence of doubly ionized ^25^Mg^2+^, which implies the absence of doubly ionized ^24^Mg and ^26^Mg.

### Glucose and rubidium uptake measured in contracting mouse muscles

Muscle contraction stimulates glucose uptake by insulin-independent mechanism(s) that have not been comprehensively described [11]. Muscle contraction also dramatically stimulates the NKA [20].

We used the multi-channel capability of ICP-MS to measure total ^13^C-[6C]-labeled-D-glucose and Rb uptake rates in the same actively contracting muscles. Rb is an excellent congener for K uptake by the NKA [19] and provides a better signal/noise than K, and specificity versus the endogenous K which is present at high concentration in mammalian tissues.

Because contraction-related signal(s) are proposed to stimulate insulin-<u>in</u>dependent glucose uptake, and because NKA activity increases the uptake of Rb, we aimed to study the co-transport of these two analytes in contracting muscles.

Muscle contraction stimulated both glucose (Fig. 4A) and Rb uptake (Fig. 4B), as expected. The contraction-related rate of glucose uptake was 10.812 µM glucose/g-min, which represents contraction-related, non-insulin-dependent glucose uptake rate. The Rb uptake rate in the same muscles was ∼1.200 µMol Rb/g-min.

### Measurement of ^3^H-glucose uptake and comparison with^13^C-glucose uptake measured by ICP-MS

To validate our measurement of ^13^C-[6C]-labeled-D-glucose uptake by ICP-MS, we measured glucose uptake using tracer ^3^H-2-deoxy-glucose. Paired EDL muscles were subjected to the same stimulation protocol but with a tracer amount of ^3^H-glucose included in the incubation Buffer. Under our experimental conditions, the glucose uptake by resting muscles was below the limit of quantification for both the ICP-MS and ^3^H measurements. The mean ^3^H-glucose uptake rate in stimulated muscles was 9,887 nMol/g-min ± 220 (n=3) which compares well with the glucose uptake rate measured by ICP-MS (11,812, Fig. 4).

### Limits of Detection

The ICP-MS based limit of detection was calculated as 3*SD of the calibration blank/slope of the calibration curve for the ^13^C isotope. In our case, two different calibrations can be used, one based on the inorganic C in water (Figure 1a), and one based on the extra-^13^C against the spiking level on the tissue to account for the endogenous background of carbon, as seen in figure 1. Under our conditions, the instrument LOD for water matrix was 0.0015 ± 0.001 ppm, while the LOD for concentrated tissue extract was 0.014 ppm ± 0.01 ppm. The LOD at the tissue level depends on the dilution factor and extraction of endogenous C. The lowest spiking intended for this method in its current form was of 0.08 ppm expressed as ^13^C, which in our current method is equivalent to 1.15 µM glucose in solution or 60-170 µM glucose in tissue, which is well below the normal mM concentration of glucose in blood or cells.

## DISCUSSION

We developed a sensitive, ratiometric method to measure ^13^C in mammalian tissues by ICP-MS, using ^13^C-[6C]-labeled-D-glucose as our test compound. The method efficiently extracts ^13^C-[6C]-labeled-D-glucose and other polar analytes from mammalian tissues and detects ^13^C with a LOD of 70-190 µM glucose in tissue for ^13^C in bovine liver CRM. While the LOD obtained cannot compete with traditional quantification of metals by ICP-MS, this level of ^13^C detection is entirely new and extends the use of ICP-MS to a wide range of biomedical research applications, where changes in the ppm range are biologically relevant. As proof-of-principle, we validated the method with measurements of glucose and Rb uptake in contracting mouse skeletal muscles *ex vivo*.

### Carbon detection by ICP-MS in biological samples

Historically, biomedical applications of ICP-MS have been limited to detection of inorganic metals and semi-metal ions such as K, Zn, Fe, As, and Se in cells and tissues, including micro-samples [14, 13]. In contrast, only limited applications have been successfully implemented for carbon-based molecules, mostly for laser ablation studies as an internal standard or quality control for acid mineralization, where residual carbon can degrade instrument performance. Recently an application to characterize and quantify micro-plastics in water was developed with ultra-fast detector mode at m/z=13, with calibration with inorganic carbon based only on the ^13^C signal[17, 18]. The use of ^13^C-enriched materials is essential for carbon analysis by ICP-MS because the endogenous contribution of organic bio-molecules makes the analysis of natural abundance carbon too variable and nonspecific to be useful. However, measurement of ^13^C by ICP-MS has not yet achieved wide use in biomedical research due to the high carbon background of biological samples as well as the high ionization energy of carbon, potential interferences from divalent ions, and difficulties in sourcing ^13^C-enriched materials.

In contrast, molecular mass spectrometry has been widely employed to detect ^13^C-labeled molecules for metabolomic and proteomic studies for over a decade. Molecular fragmentation and incorporation of liquid or gas chromatography before mass spectrometry are critical for these approaches. The main drawbacks of molecular mass spectrometry are the low ionization efficiency and matrix dependent ionization that can result in ion suppression, and the complexity of the resulting mass spectra. Because a single molecule can exist in several ionization states or associate with various common molecules (water, sodium, calcium et al.) and most molecules present in the ionization interface are reflected in the obtained spectra, the spectra are difficult to interpret without fragmentation. In addition, soft ionization methods commonly used to preserve the integrity of target molecules are sensitive to matrix overloading and signal suppression, and accurate quantification is only possible with heavy isotope spiking of individual target molecules. The use of a strong ionization source in the form of argon plasma present in an ICP-MS can reduce the dependence of results on matrix composition, simplify interpretation, and improve quantification.

With the goal of developing a simple method to detect ^13^C-based molecules for a wider range of biomedical applications, we focused our efforts on ^13^C-[6C]-labeled-D-glucose. The success of our method depended on: i) avoiding extraction of as much background natural abundance carbon from the sample as possible while extracting ^13^C-labeled glucose and target ions efficiently; and ii) finding instrument conditions for a stable ^13^C/^12^C yet responsive signal without spectral interferences.

The first goal was accomplished by disrupting the tissues with probe-assisted ultra-sonication in 18 MΩ water. Water was a successful extractant for these cytosolic, polar analytes. Ultra-sonication gave enhanced reproducibility compared to mortar-and-pestle or acid dissolution, without forming elemental carbon (charcoal). Extraction efficiency, evaluated based on recovery of a certified content of Rb and spiked ^13^C-[6C]-labeled-D-glucose, was > 80% for Rb and > 95 % for ^13^C. The extraction protocol could be refined to further reduce the endogenous background carbon, but was adequate for the test application used in this study

The instrument parameters developed for detecting ^13^C-[6C]-labeled-D-glucose by ICP-MS were guided by previous work [19, 16]. The ability to quantify ^13^C-[6C]-labeled-D-glucose requires a ^13^C/^12^C signal variation that is sufficiently above the background noise of a sample containing only natural abundance carbon. Measurement of a daily ^13^C/^12^C ratio is required because the sensitivity of an ICP-MS instrument for these ions depends on the inherent mass bias of extraction cones, ion lenses, collision/reaction cells, mass filters and detectors. The use of a high power, 1600-watt plasma torch under no-gas tune, with the collision/reaction cell in bypass mode, improved the stability of the ^13^C/^12^C ratio. The reduction of ion-beam differential deflection in the omega bias/omega lenses as well as the Oct bias/QP bias was necessary for a stable ^13^C/^12^C ratio over a wide range of C content. Despite day-to-day variability, the instrument ^13^C/^12^C ratio was stable within reported values (<3%) for the duration of a six-hour run, and allowed analysis of well over a hundred samples with minimal drift.

The use of the isotopic content of both ^12^C and ^13^C was necessary as the standard addition method would rely on the same background of C, something not achievable by our method. By obtaining an instrumental ^13^C/^12^C ratio from control tissues, the calculated extra-^13^C CPS were used to calculate the extra ^13^C concentration. This calculation is only valid because the sensitivity of the instrument to each carbon isotope was the same regardless of the sample matrix, carbon source or variabilities in the endogenous C background. The reproducibility of the calibrations, natural abundance-instrumental ^13^C/^12^C ratio and instrumental optimizations allowed for a robust quantification of ^13^C-[6C]-labeled-D-glucose in contracting muscles. The LOD in µM are well suited for this analysis, given that the extra and intra cellular concentrations of glucose and its metabolites in muscle are in the milli-molar range.

Although a remote possibility, we examined whether ^24^Mg^2+^ and ^26^Mg^2+^ might be present as interferences at the same m/z as ^12^C and ^13^C, respectively. This was an important part of our method development due to the high content of Mg in biological samples and the low levels of detection required for target elements. If ^24^Mg^2+^ and ^26^Mg^2+^ signals were to add to the signals, a complex mathematical equation would have been needed to identify the carbon signal. This correction would add variability in proportion to the Mg concentration and sample composition, and negatively affect the LOD. Our results demonstrate that the ^13^C/^12^C signal is <u>independent</u> of Mg concentration or sample composition and depends linearly on the concentration of ^13^C-[6C]-labeled-D-glucose in the spiked samples.

### Glucose uptake in contracting mouse skeletal muscle measured by ICP-MS

As proof-of-principle, we measured glucose uptake in contracting mouse EDL muscles, which have a highly glycolytic metabolism. We chose this experimental model because glucose uptake increases greatly, in the milli-molar range, during muscle contraction and because the skeletal muscles play a major role in glucose homeostasis. Glucose uptake in quiescent muscles was much lower than the contracted pair; this result was expected because basal glucose uptake requires insulin from the circulation, which is negligible in the ex vivo model. Contraction dramatically stimulated glucose uptake, as expected. The contraction-related rate of glucose uptake measured by ICP-MS compared well with that measured using a conventional radiolabeled tracer glucose assay, which further validates the method

In addition, we combined the ^13^C sensitivity of our method with the multi-element capability of ICP-MS to measure glucose and Rb uptake in the same contracting muscles. This application was motivated by the fact that glucose uptake by contracting muscles is not completely understood and. In contrast to basal glucose uptake, contracting muscles take up glucose without a requirement for insulin. Indeed, circulating insulin declines during prolonged or intense exercise. A number of studies have proposed mechanisms by which GLUT4 translocation may be triggered by signaling pathways that do not require the Insulin Receptor [11](reviewed in Richter & Hargreaves 2015;). Notably, a large fraction of glucose uptake persists in the absence of GLUT4 (GLUT4 KO), suggesting the existence of an additional glucose transporter(s) in muscle.

A more complete characterization of alternative signaling pathways for GLUT4 translocation, as well as the identification of glucose uptake mechanisms other than GLUT4, will require experimental approaches that allow simultaneous measurement of glucose and multiple other factors or ions. The ^13^C sensitivity of our method and the multi-element capability of ICP-MS are well-suited for this application. To evaluate this approach, we measured glucose uptake simultaneously with K/Rb uptake by the NKA in contracting muscles. This test was chosen because NKA activity, specifically NKA α2 isoform activity, increases dramatically during muscle contraction [21]; and because NKA activity generates the Na gradient that provides the driving force for many other transport processes. Rb uptake increased due to contraction-related stimulation of the NKA, as expected.

### Biological applications and extensions of the method

Our method for detecting ^13^C-enriched molecules together with inorganic ions has wide applications for biomedical research. The method developed for ^13^C-labeled glucose can be further developed and broadly applied to detect other organic molecules and metabolites where changes in the µM range are biologically relevant. The multi-channel capability of ICP-MS is a particular advantage for studies of coupled or secondary transport mechanisms that involve multiple ions species and/or transporters. These applications include many vital cotransporters and exchangers (Na/Ca exchanger, anion exchangers et al.) or secondary active transporters (SGLTs, sodium-coupled nutrient transporters, neurotransmitter transporters et al.) that use the energy stored in the Na gradient generated by NKA activity to drive uptake of essential ions and nutrients into cells and organelles. This capability can also be applied to studies of complex molecular interactions in other signaling and metabolic pathways.

## Conclusion

In conclusion, we successfully developed, validated and applied a method, based on water-based ultra-sonication extraction to quantitatively extract Rb and glucose from whole muscle tissues. The extraction procedure was suitable for ICP-MS detection, and for this, we developed a method to quantify ^13^C-[6C]-labeled-D-glucose and Rb simultaneously. The developed approach was applied to an *ex-vivo* electrically-stimulated, muscle contraction-induced glucose uptake model with results comparable to an established method in the form of ^3^H-2-DG uptake. The safety advantages over radioisotope tracer measurements and the multi-ion flux detection capabilities of ICP-MS bring our method to the toolbox of researchers interested in the study of complex flux analysis and other complex molecular interactions.

## ACKNOWLEDGMENTS

Part of this work was funded by the NIH Grant RO1 AR063710.

## FIGURES

**Supplemental Figure 1**

Example of one set of calibration curves in the original format obtained in the Agilent Mass Hunter software, showing calibration data details for inorganic carbon standard based ^13^C and (b) ^12^C and (c) ^13^C generated with ^13^C-[6C]-labeled-D-glucose.

**Supplemental Figure 2**

Rubidium response in CPS for the magnesium spiking experiment. (a) carbon, magnesium and carbon + magnesium in water and (b) in bovine liver CRM extracts. The scale is log2 to cover the broad range of instrument response. The numbers on top of each bar represent the spiking level of carbon, magnesium or carbon + magnesium. Error bars represent the standard deviation from 3 experimental replicates.

**Supplemental Figure 3**

Continuous-line representation of the mass spectra obtained from water and water spiked with ^13^C-[6C]-labeled-D-glucose (black continuous and black segmented line); and bovine liver CRM extracts and spiked bovine liver CRM extracts (gray continuous and gray segmented line). The arrows represent the gain in count rate at m/z=13 on both spiking experiments. The acquisition was in semi-quantitative mode to capture 10 points per m/z unit.

