## Supplementary figures and images for "Glucose uptake in mammalian cells measured by ICP-MS"

### Supplemental figure 1

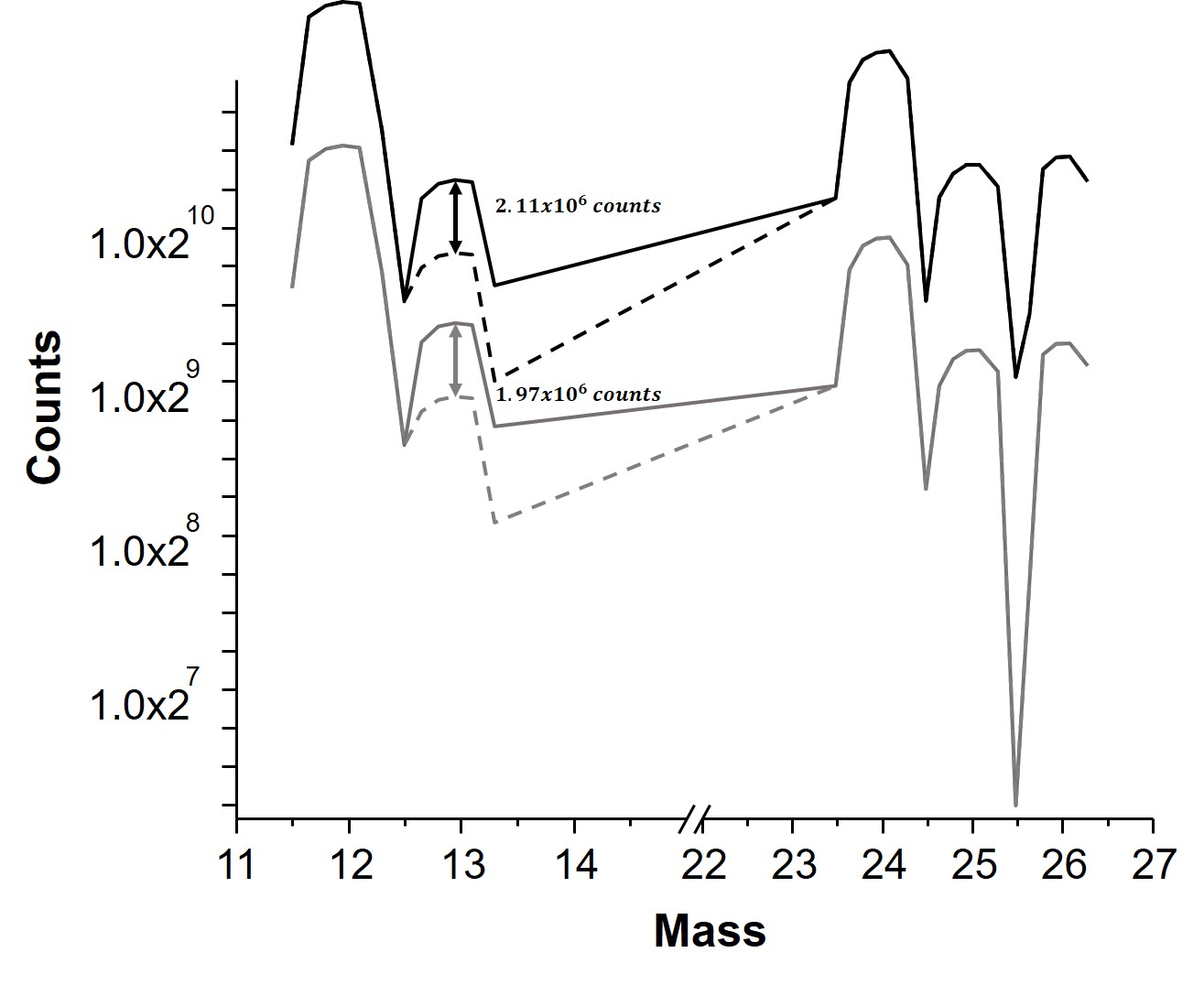

### Supplemental figure 2

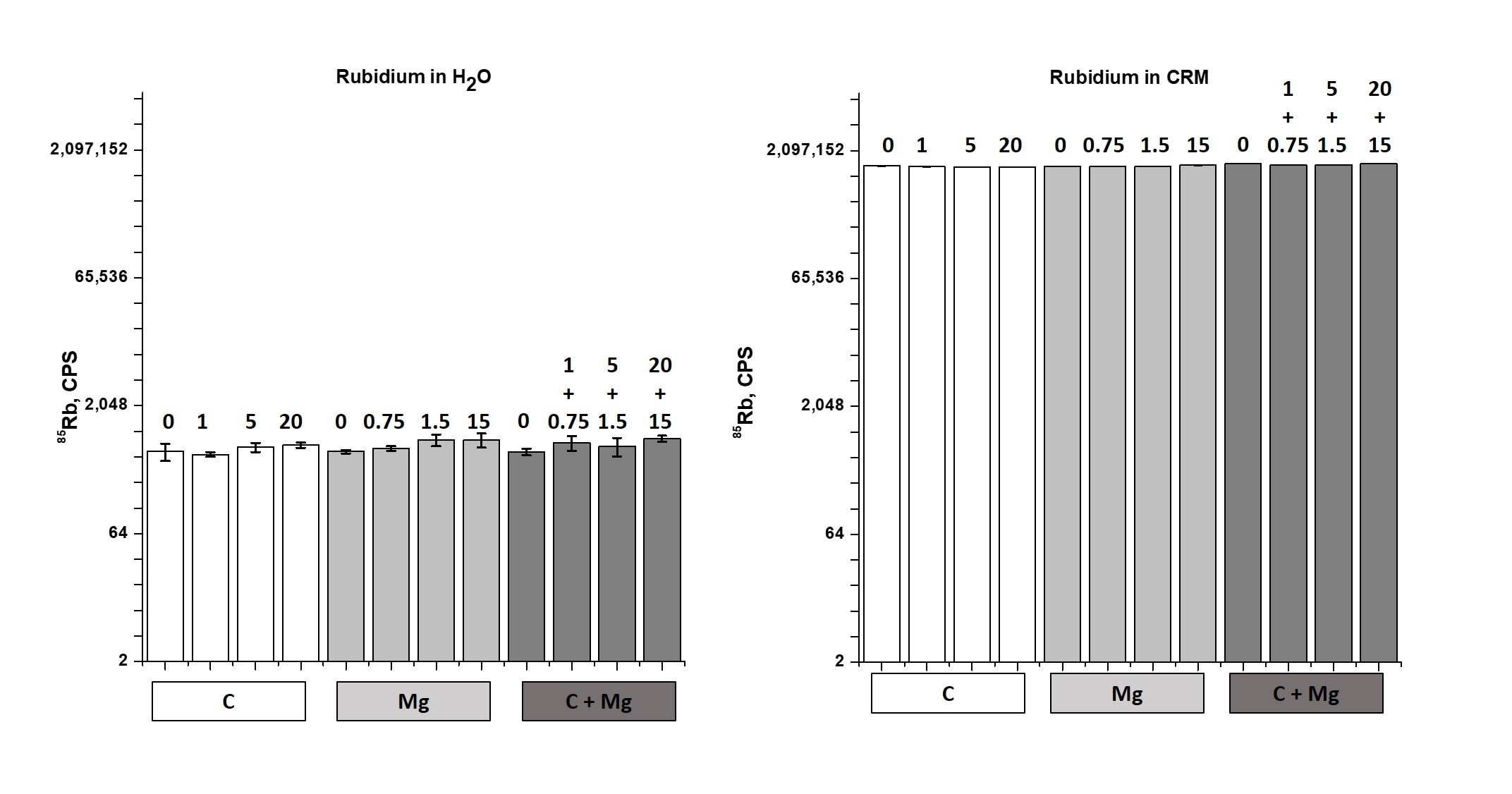

### Supplemental figure 3

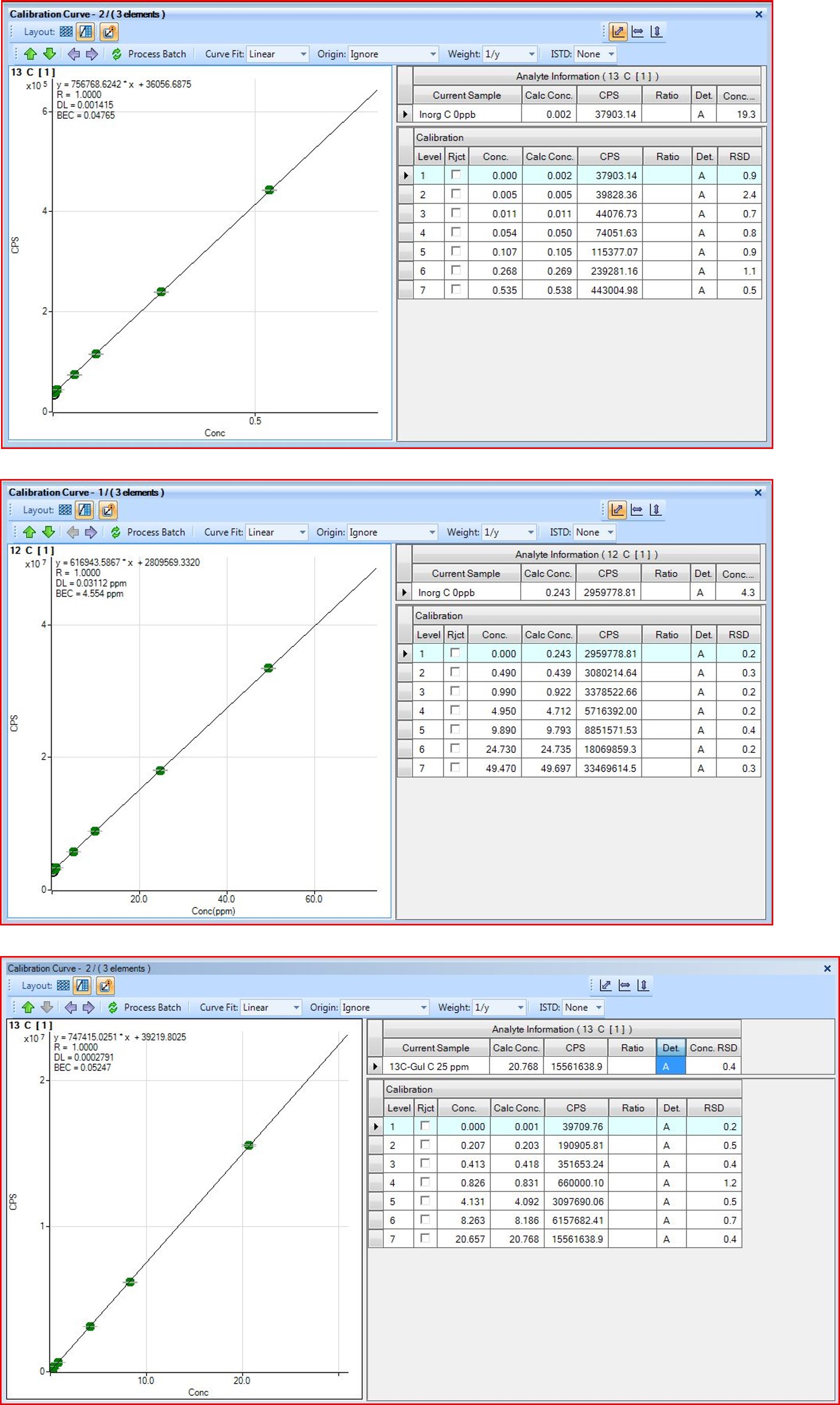
